# Somatic aberrations of BRCA1 gene are associated with progressive and stem cell-like phenotype of prostate cancer

**DOI:** 10.1101/271312

**Authors:** Aleksandra Omari, Paulina Nastały, Aneta Bałabas, Michalina Dąbrowska, Beata Bielińska, Sebastian Huss, Klaus Pantel, Axel Semjonow, Elke Eltze, Burkhard Brandt, Natalia Bednarz-Knoll

**Author notes:** **Corresponding author:** Dr Natalia Bednarz-Knoll, Laboratory of Cell Biology, Department of Medical Biotechnology, Medical University of Gdańsk, Gdańsk, Poland; formerly Institute of Tumour Biology, University Medical Centre Hamburg-Eppendorf, Hamburg, Germany.

## Abstract

**Background:** BRCA1 is a pivotal tumor suppressor. Its dysfunction is known to play a role in different tumor entities. Among others, BRCA1 germline mutations account for higher risk and more aggressive course of prostate cancer (PCa). In addition, somatic BRCA1 gene loss was demonstrated to be a signature of PCa dissemination to regional lymph nodes and peripheral blood, and indicate worse clinical outcome. In order to substantiate the data for BRCA1 gene loss in PCa and to reveal its phenotypical background, BRCA1 gene status was assessed in a large cohort of PCa patients and compared to different molecular factors.

**Methods:** BRCA1 gene dosage was assessed in 2398 tumor samples from 1199 PCa patients using fluorescent in situ hybridization. It was compared to clinic-pathological parameters, patients’ outcome as well as selected proteins (Ki-67, apoptosis marker, cytokeratins, vimentin, E- and N-cadherin, ALDH1 and EGFR) examined by immunohistcohemistry.

**Results:** BRCA1 losses were found in 10%, whereas gains appeared in 7% of 603 informative PCa patients. BRCA1 losses correlated to higher T status (p=0.027), Gleason score (p=0.039), shorter time to biochemical recurrence in patients with Gleason score >7 independently of other factors (multivariate analysis, p=0.005) as well as expression of proteins regulating stemness and epithelial-mesenchymal transition i.e. ALDH1 (p=0.021) and EGFR (p=0.011), respectively. BRCA1 gains correlated to shorter time to metastasis (p=0.012) and expression of ALDH1 (p=0.014).

**Conclusions:** The presented results support the assumption that BRCA1 gene losses contribute to a progressive and stem cell-like phenotype of PCa. Furthermore, they reveal that also BRCA1 gain might mark more invasive tumors.

## Background

Previous analysis of primary prostate cancer (PCa), its metastasis to lymph nodes and circulating tumor cells (CTCs) revealed that loss of the prominent tumor suppressor gene BRCA1 can be one signature of PCa aggressiveness and its dissemination to regional lymph nodes and peripheral blood [1]. BRCA1 function-off-mutations are well described in development and progression of different solid tumors particularly breast and ovarian cancers but also PCa. Germline BRCA1 mutations are shown to be associated with the higher risk of the onset of PCa and/or more aggressive course of the disease [2–4]. Of note, BRCA1 dysfunction might be particularly interesting in the context of poly(ADP-ribose) polymerases (PARP) inhibitor-based therapies, tested currently in some clinical trials including also patients with PCa in such trials as TOPARP trial (NCT01682772), PROFound study (NCT02987543) and TRITON3 (NCT02975934) [5]. The identified gains of BRCA1 seemed to be irrelevant for the PCa patients’ outcome but it might have been biased by the low number of cases with such characteristics [1].

The molecular associations of BRCA1 gene loss in PCa remain unclear. BRCA1 is known to be a pivotal regulator of a plethora of cellular processes, which, when defected, might be involved in tumor progression [6]. It includes cell cycle control, DNA repair, transcription and ubiquitination [7–8]. BRCA1-mutated or -deficient breast cancer cells seem to have more invasive phenotype including higher proliferation rate [9[ and migration abilities [10–12]. Moreover, in breast cancer *in vitro* and mouse models it was also shown that BRCA1 alterations could result in epithelial-mesenchymal transition (EMT) [10,13,14] and even influence cellular fate by inducing expression of proteins involved in stemness programme [15,16]. Of note, in peripheral blood of metastatic PCa patients, it seemed that mesenchymal cell marker vimentin identified cells with BRCA1 loss more frequently than common cytokeratin (CK) staining [1], which might suggest that such a phenomenon takes place also in PCa.

In the current work, the clinical associations of BRCA1 gene aberrations found in the previous studies [1,17] were evaluated on the large cohort of patients. Additionally, to investigate potential phenotype of BRCA1-gene-deficient PCa, expression of selected proteins in PCa was determined in relation to the status of BRCA1 gene. The set of the analysed proteins included proliferation (Ki-67) and apoptosis markers (ApopTags), basal (CK 5/6, CK14) and luminal cytokeratins (CK8/18 and CK19), mesenchymal marker vimentin, two cadherins (E- and N- cadherin) which changes are known to be associated with EMT induction, stem cell marker (ALDH1) as well as potent regulator of cellular growth, stemness, and EMT (EGFR).

## Methods

### Patients

One thousand one hundred ninety nine patients who underwent radical prostatectomy at the Department of Urology University Hospital Münster (Germany) during 1993-2004 were included in the current study. The patients were proven to suffer from mono- or multifocal PCa. The tumors were characterized by different clinical parameters such as TNM status (according to AJCC Cancer Staging 2010) or Gleason score (Tab. 1). The mean patient age at the moment of surgery was 64 years old (the range: 46 - 81), and pathologic stages of their tumours ranged from pT2pN0cM0 to pT4pN1cM0. The patients’ disease free (DFS), metastasis free (MFS) and overall survival (OS) data were collected. Biochemical recurrence was defined as two consecutive concentrations of prostate specific antigen (PSA) above 0.1 ng/mL The time point of recurrence was defined as the first PSA concentration above 0.1 ng/mL The last follow-up was performed in October 2014. In average, the patients were observed for 60 months. The maximum observation period lasted 223 months (approx. 18 years). Information about patients’ death and occurrence of overt metastasis was documented. The study was conducted according to REMARK study recommendations [18].

**Table 1.** Associations of BRCA1 gene aberrations to clinical data. Comparison of BRCA1 gene status to clinico-pathological parameters with the usage of Fisher exact test (F) or otherwise Chi square test. Neg indicates negative expression, pos – positive expression, BR - biochemical relapse, L – gene loss, N – normal gene dosage, G – gene gain. Note that not all numbers sum up to 1199 due to the missing data.

| BRCA1 gene dosage |  |  |  |  |  |  |  |  |  |  |
| --- | --- | --- | --- | --- | --- | --- | --- | --- | --- | --- |
|  | Patients |  | Loss |  | Normal |  | Gain |  | p-values |  |
|  | n | % | n | % | n | % | n | % | L vs. N | G vs. N |
| **Age** |  |  |  |  |  |  |  |  |  |  |
| <64 | 623 | 52.0 | 28 | 45.9 | 259 | 51.7 | 18 | 43.9 |  |  |
| >64 | 574 | 48.0 | 33 | 54.1 | 242 | 48.3 | 23 | 56.1 |  |  |
| **total** | **1997** |  | **603** |  |  |  |  |  | **0.393** | **0.337** |
| **T status** |  |  |  |  |  |  |  |  |  |  |
| T2 | 504 | 42.6 | 14 | 23.0 | 187 | 38.1 | 10 | 24.4 |  |  |
| T3 | 635 | 53.7 | 46 | 75.4 | 283 | 57.6 | 31 | 75.6 |  |  |
| T4 | 43 | 3.6 | 1 | 1.6 | 21 | 4.3 | 0 | 0 |  |  |
| **total** | **1182** |  | **593** |  |  |  |  |  | **0.027** | **0.057** |
| **N status** |  |  |  |  |  |  |  |  |  |  |
| N0 | 1086 | 92.7 | 52 | 88.1 | 452 | 92.1 | 36 | 90.0 |  |  |
| N1 | 86 | 7.3 | 7 | 11.9 | 39 | 7.9 | 4 | 10.0 |  |  |
| **total** | **1172** |  | **590** |  |  |  |  |  | **0.304** | **0.646** |
| **Gleason Grading Score** |  |  |  |  |  |  |  |  |  |  |
| <7 | 343 |  | 6 | 10.0 | 117 | 23.4 | 5 | 12.2 |  |  |
| 3+4 | 369 |  | 16 | 26.7 | 155 | 31.0 | 14 | 34.1 |  |  |
| 4+3 | 301 |  | 22 | 36.7 | 138 | 27.6 | 13 | 31.7 |  |  |
| >7 | 185 |  | 16 | 26.7 | 90 | 18.0 | 9 | 22.0 |  |  |
| **total** | **1198** |  | **601** |  |  |  |  |  | **0.039** | **0.426** |
| **Focality** |  |  |  |  |  |  |  |  |  |  |
| uni | 152 | 13.2 | 6 | 9.8 | 67 | 13.6 | 3 | 7.3 |  |  |
| multi | 998 | 86.7 | 55 | 90.2 | 427 | 86.4 | 38 | 92.7 |  |  |
| **total** | **1150** |  | **596** |  |  |  |  |  | **0.547 (F)** | **0.338 (F)** |
| **BR** |  |  |  |  |  |  |  |  |  |  |
| no | 702 | 72.7 | 25 | 55.6 | 271 | 69.0 | 19 | 61.3 |  |  |
| yes | 264 | 27.3 | 20 | 44.4 | 122 | 31.0 | 12 | 38.7 |  |  |
| **total** | **966** |  | **49** |  |  |  |  |  | **0.069** | **0.377** |
| **Metastatic relapse** |  |  |  |  |  |  |  |  |  |  |
| no | 1074 | 96.1 | 55 | 96.5 | 449 | 96.4 | 33 | 86.8 |  |  |
| yes | 40 | 3.6 | 2 | 3.5 | 17 | 3.6 | 5 | 13.2 |  |  |
| **total** | **1114** |  | **595** |  |  |  |  |  | **1.000 (F)** | **0.019 (F)** |
| **Death** |  |  |  |  |  |  |  |  |  |  |
| alive | 1032 | 91.0 | 51 | 98.1 | 434 | 97.7 | 31 | 91.2 |  |  |
| PCa-related | 17 | 1.5 | 1 | 1.9 | 10 | 2.3 | 3 | 8.8 |  |  |
| not Pca-related | 50 | 4.4 | - | - | - | - | - | - |  |  |
| unknown | 34 | 3.0 | - | - | - | - | - | - |  |  |
| **total** | **1134** |  | **555** |  |  |  |  |  | **1.000 (F)** | **0.057 (F)** |

### Clinical material

Clinical material was prepared as six tissue microarrays (TMAs) as described [1]. Briefly, 4-μm-thick sections of TMAs comprised of 398 tissue cores (diameter of 0.6 mm) corresponding to 199 patients each were prepared for both fluorescent *in situ* hybridization (FISH) and immunohistochemistry (IHC). The patients with multifocal PCa were represented on TMA by tissue cores obtained from two different tumor foci. Normal prostate tissues were included in TMAs and evaluated as internal controls.

### Fluorescent In Situ Hybridization for assessment of BRCA1 gene dosage

Fluorescent *in situ* hybridization (FISH) protocol was applied as described [1]. Based on the previously assessed clinical impact of BRCA1 gene [1], the cut-off to categorize BRCA1 gene dosage was defined as ≤0.75 for BRCA1 losses, 0.75-1.25 for normal gene dosage and ≥1.25 for gains. Matching lymph node metastases were evaluated for BRCA1 gene dosage before [1].

### Immunohistochemistry for BRCA1 and selected proteins

BRCA1 protein expression was performed as described (19). Briefly, deparaffinised specimens were incubated after antigen retrieval (citrate buffer, pH 6.0, 20 min. at 95C) with monoclonal mouse IgG1 anti-BRCA1 (clone MS110, Calbiochem) diluted 1:150 and developed with Dako REAL™ Detection System, Peroxidase/DAB+, Rabbit/Mouse (Dako). Localization (nuclear or cytoplasmic), intensity of the staining and percentage of the cells stained for BRCA1 were documented. Negative BRCA1 staining was determined as no or weak, whereas positive BRCA1 staining as moderate to strong staining.

Immunohistochemistry for CK5/6, CK14, CK8-18, CK19, E-cadherin, N-cadherin, vimentin, apoptotic marker, Ki-67, ALDH1 and EGFR were performed as described **Supplementary Data**.

### BRCA1 mutation analysis

BRCA1 mutations were analysed by denaturation high-performance liquid chromatography (DHPLC), carried out on an automated DHPLC instrumentation (Transgenomic, Omaha, NE). Briefly, PCR products were eluted from DNASep column (Transgenomics) with triethylammonium acetate (TEAA) buffer and acetonitrile linear gradient (54.3–63.3%) at 0.9 ml/min flow rate. The gradient and temperature required for separation of DNA hetero- and homoduplexes was determined for each exon using the Wavemaker software (Transgenomic). The separation conditions used for each exon of the BRCA1 gene are listed in Suppl. Tab. 1.

### Statistics

Statistical analysis was performed with the usage of SPSS software version 24.0 licensed for University of Gdańsk. Chi-square or Fisher’s exact tests were used in order to compare the results to molecular factors or clinico-pathological parameters. Associations between BRCA1 gene status and time to biochemical recurrence or to metastasis were evaluated using Log Rank (Mantel Cox) test and Kaplan-Meier plot. To estimate hazard risk, Cox-Hazard-Potential regression analysis (CI 95%) was done. All results were considered statistically significant if p<0.05 and highly statistically significant if p<0.001. Cases with missing data were excluded from analysis.

## Results

### Frequency of BRCA1 gene aberrations in primary prostate cancer

Normal gene dosage of BRCA1 gene was observed in normal prostate cells and its stroma. Eight-hundred-forty tumor samples from 603 patients with mono- and multifocal disease were informative for BRCA1 gene dosage. The remaining cases were excluded from the further analysis due to the insufficient number of tumor cells, overlapping of tumor cells’ nuclei or inappropriate quality of FISH signals. The average number of tumor cells analyzed per tumor sample (diameter of 0.6 mm and different tumor cell content ranging from 10 to approx. 1000) was 33 with the range of 20 to 76. Sixty-nine (8.2%) samples from 61 (10.1%) patients were characterized by the loss of BRCA1 gene with the median gene dosage of 0.59 (range 0.49 to 0.75). Normal BRCA1 copy number was found in 724 (86.2%) tumor samples from 501 (83.1%) patients. The median normal BRCA1 gene dosage was 1.00. The gains of BRCA1 were detected in 47 (5.6%) tumor samples from 41 (6.8%) patients with the median gene dosage of 1.32 (range 1.25 to 2.00). The most frequent BRCA1 copy counts in this group were 3 and 4, however, in the rare single cells even up to 12 BRCA1 signals were detected. Eleven patients characterized by BRCA1 gain were tested for 6 germline mutations of BRCA1 gene frequently found in European population: 185delAG, Cys61Gly, 3819del5, 4153delA, 5382insC and Arg1751Ter. No mutations were found in these patients (data not shown). Additionally, no BRCA1 promoter methylation was found in these patients as described [1]. When BRCA1 gene dosage and protein expression were compared in subpopulation of PCa (n=203), 4 out of 10 (40%) tumors with BRCA1 gene gain were characterized by loss of nuclear BRCA1 protein expression or its localisation in cytoplasm (Suppl. Table 2).

### Intratumoral heterogeneity of PCa in regard to BRCA1 aberrations

Two-hundred-thirty-four patients were informative for BRCA1 gene dosage in two tissue samples originated from two separated tumor foci. Both gene losses and gains were found predominantly in exclusively one tumor focus. In total, differences in BRCA1 gene dosage between two analyzed tumor samples occurred in 34 (14.5%) cases. Fourteen and 5 patients carried BRCA1 loss and gain in both tumor foci, respectively, whereas 13 and 10 patients displayed BRCA1 loss and gain exclusively in one analyzed tumor focus. Interestingly, one patient had BRCA1 loss in one focus and gain in another one (Suppl. Tab. 3).

### Frequency of BRCA1 gene aberrations in matched pairs of prostate cancer - lymph node metastases

Thirty patients were analysed for matched primary tumor (PT) – lymph node metastasis (LNM) pairs based on the current and previous results [1]. Eleven (36.7%) of 30 patients revealed loss in LNMs. BRCA1 loss was detected in both primary tumor and LNMs in 4 (13.3%) patients (Suppl. Tab. 4). No patients revealed gains in LNM.

### BRCA1 gene aberrations are associated with poor clinical outcome

Both BRCA1 gene losses occurred more frequently in patients with more advanced tumors (Chi^2^=7.236, p=0.027) (Tab. 1). In addition, BRCA1 gene losses correlated to more advanced Gleason score (Chi^2^=8.376, p=0.039), whereas BRCA1 gene gains were significantly associated with metastatic relapse (Fisher exact test, p=0.019) (Tab. 1).

In the total cohort of patients, BRCA1 gene gains predicted shorter time to metastasis (Chi^2^=6.312, p=0.012) (Fig. 1A). In the patients characterized by Gleason score >7, BRCA1 losses correlated with shorter time to biochemical recurrence (Chi^2^=5.280, p=0.022; median time to biochemical recurrence 27 vs. 113 months for patients with BRCA1 gene loss or normal gene dosage, respectively) (Fig. 1B). In this subgroup of patients, BRCA1 gene losses appeared also to be an independent prognostic marker of shorter time to biochemical recurrence (Cox analysis, p=0.005, CI95% 0.336, HR 0.158-0.715) (Tab. 2).

**Figure 1.**
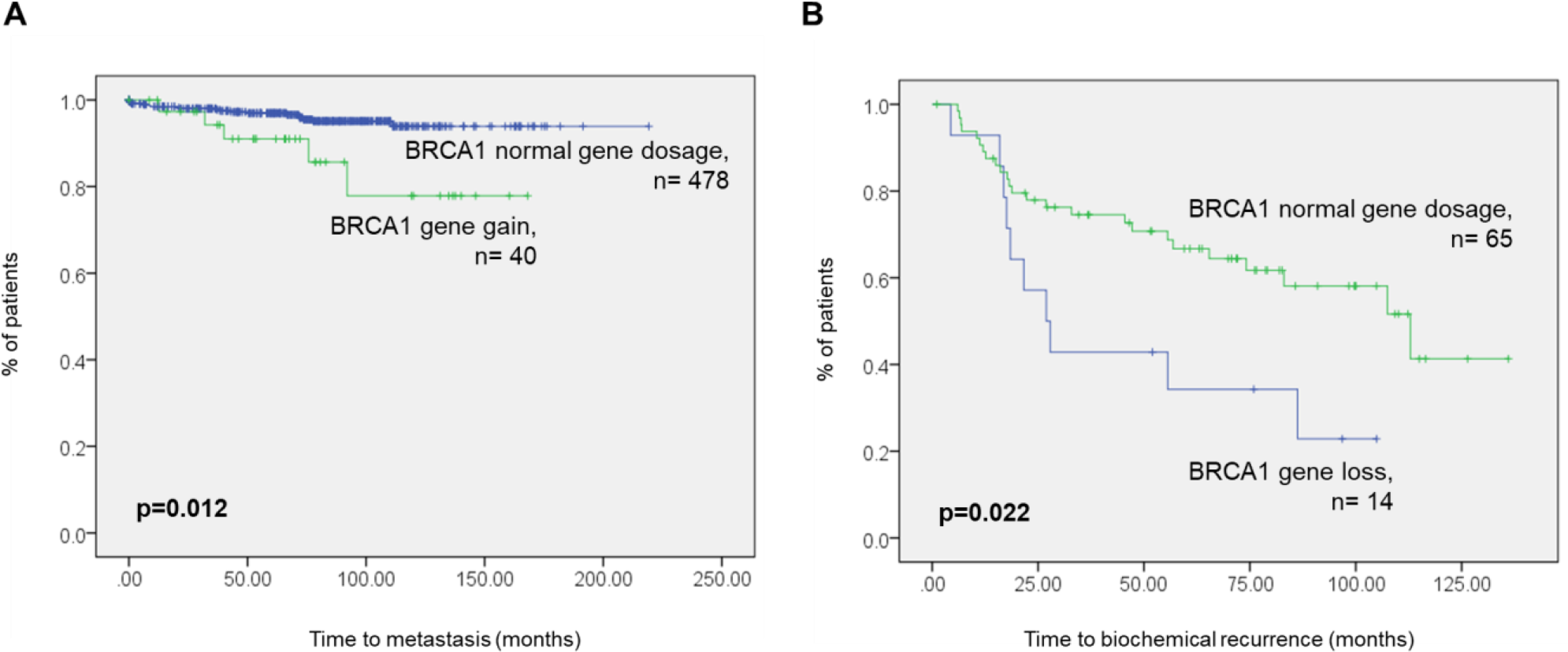
Survival analysis. Associations of BRCA1 gene gains to shorter time to metastasis in total cohort of patients (A) and BRCA1 gene losses to shorter time to biochemical recurrence in patients with Gleason score >7 (B).

**Table 2.**
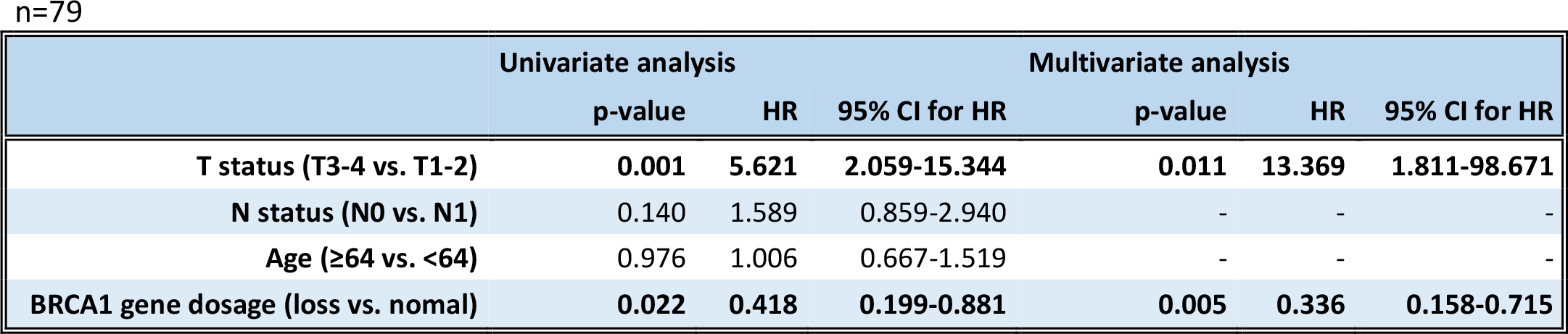
Multivariate analysis.

### Associations of BRCA1 gene aberrations with cancer phenotypes

Primary tumors with BRCA1 gene losses and gains displayed significantly more frequently stem cell marker ALDH1 (Fisher exact test, p=0.021 and p=0.014, respectively) (Tab. 3). However, there was no correlation of BRCA1 gene status to ALDH1 staining in LNMs (data not shown). BRCA1 gene losses showed also a correlation to the expression of EGFR (Fisher exact test, p=0.011) (Tab. 3). None of BRCA1 aberrations were associated with proliferation status, events of apoptosis or changes in the pattern of epithelial (E-cadherin, cytokeratins) and mesenchymal markers (N-cadherin and vimentin) (Tab. 3).

**Table 3.**
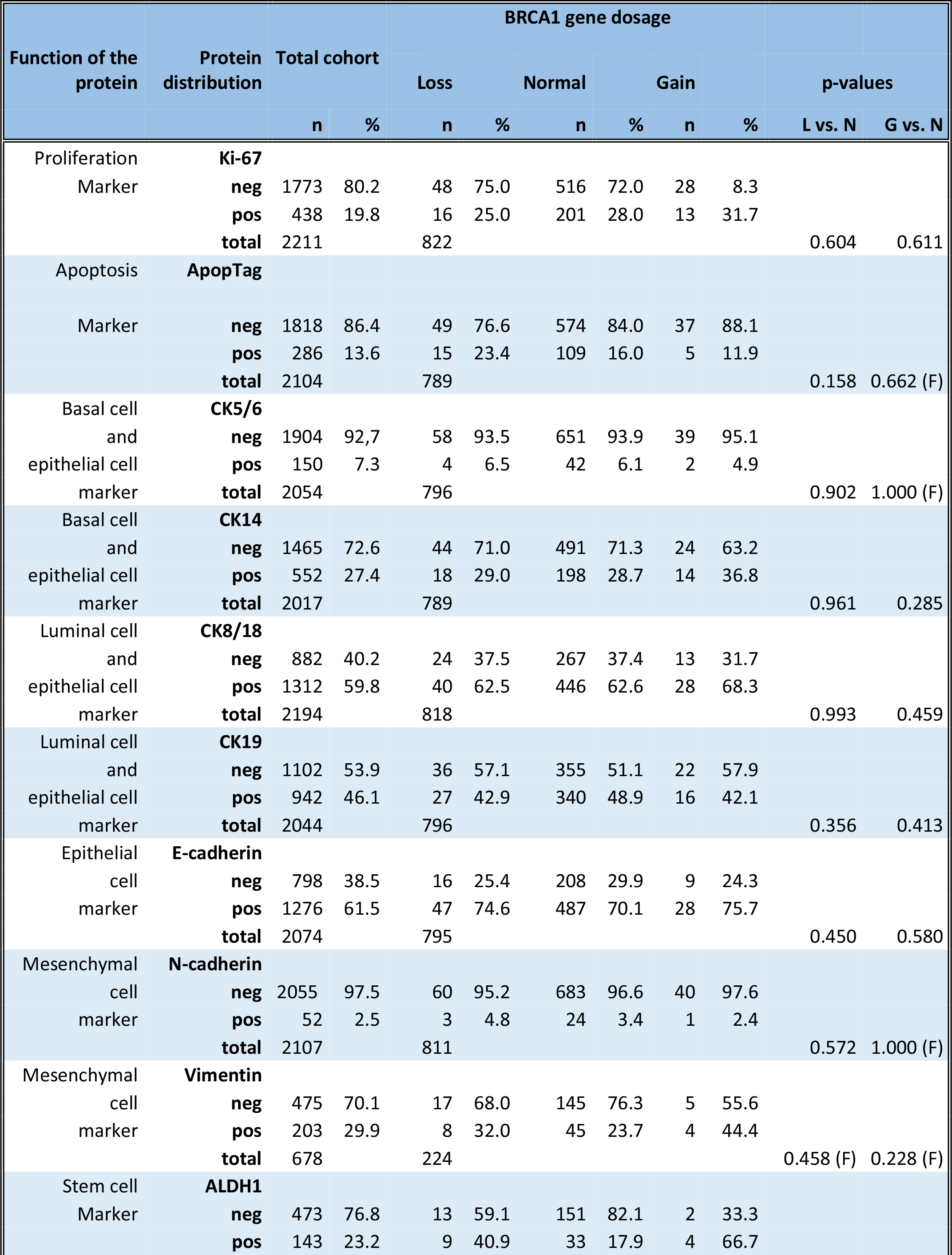
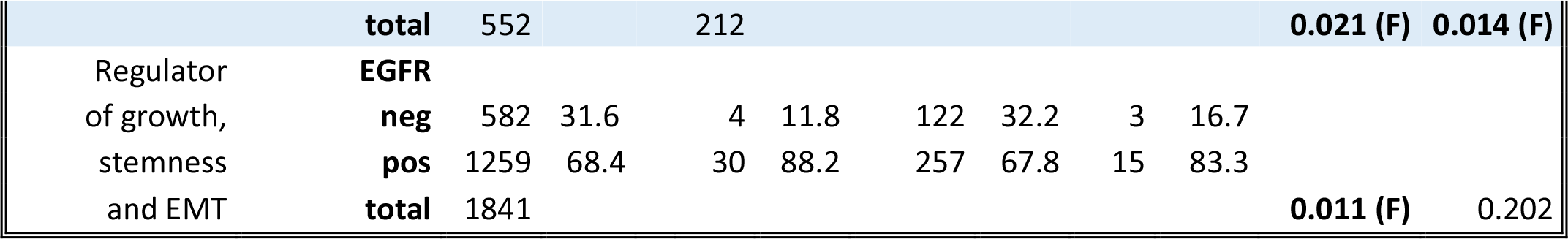
Associations of BRCA1 gene aberrations to molecular data. Comparison between BRCA1 gene status and immunohistochemically assessed proteins was done in tissue cores (n=796 for vimentin and ALDH1, whereas n=2398 for all other markers) with the usage of Fisher exact test (F) or otherwise Chi square test. Neg indicates negative expression, pos – positive expression, L – gene loss, N – normal gene dosage, G – gene gain.

| BRCA1 gene dosage |  |  |  |  |  |  |  |  |  |  |  |
| --- | --- | --- | --- | --- | --- | --- | --- | --- | --- | --- | --- |
| Function of the protein | Protein distribution | Total cohort |  | Loss |  | Normal |  | Gain |  | p-values |  |
|  |  | n | % | n | % | n | % | n | % | L vs. N | G vs. N |
| Proliferation Marker | Ki-67 |  |  |  |  |  |  |  |  |  |  |
|  | neg | 1773 | 80.2 | 48 | 75.0 | 516 | 72.0 | 28 | 8.3 |  |  |
|  | pos | 438 | 19.8 | 16 | 25.0 | 201 | 28.0 | 13 | 31.7 |  |  |
|  | total | 2211 |  | 822 |  |  |  |  |  | 0.604 | 0.611 |
| Apoptosis | ApopTag |  |  |  |  |  |  |  |  |  |  |
|  | neg | 1818 | 86.4 | 49 | 76.6 | 574 | 84.0 | 37 | 88.1 |  |  |
|  | pos | 286 | 13.6 | 15 | 23.4 | 109 | 16.0 | 5 | 11.9 |  |  |
|  | total | 2104 |  | 789 |  |  |  |  |  | 0.158 | 0.662 (F) |
| Basal cell and epithelial cell marker | CK5/6 |  |  |  |  |  |  |  |  |  |  |
|  | neg | 1904 | 92.7 | 58 | 93.5 | 651 | 93.9 | 39 | 95.1 |  |  |
|  | pos | 150 | 7.3 | 4 | 6.5 | 42 | 6.1 | 2 | 4.9 |  |  |
|  | total | 2054 |  | 796 |  |  |  |  |  | 0.902 | 1.000 (F) |
| Basal cell and epithelial cell marker | CK14 |  |  |  |  |  |  |  |  |  |  |
|  | neg | 1465 | 72.6 | 44 | 71.0 | 491 | 71.3 | 24 | 63.2 |  |  |
|  | pos | 552 | 27.4 | 18 | 29.0 | 198 | 28.7 | 14 | 36.8 |  |  |
|  | total | 2017 |  | 789 |  |  |  |  |  | 0.961 | 0.285 |
| Luminal cell and epithelial cell marker | CK8/18 |  |  |  |  |  |  |  |  |  |  |
|  | neg | 882 | 40.2 | 24 | 37.5 | 267 | 37.4 | 13 | 31.7 |  |  |
|  | pos | 1312 | 59.8 | 40 | 62.5 | 446 | 62.6 | 28 | 68.3 |  |  |
|  | total | 2194 |  | 818 |  |  |  |  |  | 0.993 | 0.459 |
| Luminal cell and epithelial cell marker | CK19 |  |  |  |  |  |  |  |  |  |  |
|  | neg | 1102 | 53.9 | 36 | 57.1 | 355 | 51.1 | 22 | 57.9 |  |  |
|  | pos | 942 | 46.1 | 27 | 42.9 | 340 | 48.9 | 16 | 42.1 |  |  |
|  | total | 2044 |  | 796 |  |  |  |  |  | 0.356 | 0.413 |
| Epithelial cell marker | E-cadherin |  |  |  |  |  |  |  |  |  |  |
|  | neg | 798 | 38.5 | 16 | 25.4 | 208 | 29.9 | 9 | 24.3 |  |  |
|  | pos | 1276 | 61.5 | 47 | 74.6 | 487 | 70.1 | 28 | 75.7 |  |  |
|  | total | 2074 |  | 795 |  |  |  |  |  | 0.450 | 0.580 |
| Mesenchymal cell marker | N-cadherin |  |  |  |  |  |  |  |  |  |  |
|  | neg | 2055 | 97.5 | 60 | 95.2 | 683 | 96.6 | 40 | 97.6 |  |  |
|  | pos | 52 | 2.5 | 3 | 4.8 | 24 | 3.4 | 1 | 2.4 |  |  |
|  | total | 2107 |  | 811 |  |  |  |  |  | 0.572 | 1.000 (F) |
| Mesenchymal cell marker | Vimentin |  |  |  |  |  |  |  |  |  |  |
|  | neg | 475 | 70.1 | 17 | 68.0 | 145 | 76.3 | 5 | 55.6 |  |  |
|  | pos | 203 | 29.9 | 8 | 32.0 | 45 | 23.7 | 4 | 44.4 |  |  |
|  | total | 678 |  | 224 |  |  |  |  |  | 0.458 (F) | 0.228 (F) |
| Stem cell Marker | ALDH1 |  |  |  |  |  |  |  |  |  |  |
|  | neg | 473 | 76.8 | 13 | 59.1 | 151 | 82.1 | 2 | 33.3 |  |  |
|  | pos | 143 | 23.2 | 9 | 40.9 | 33 | 17.9 | 4 | 66.7 |  |  |
|  | **total** | 552 |  | 212 |  |  |  |  |  | **0.021 (F)** | **0.014 (F)** |
| Regulator | **EGFR** |  |  |  |  |  |  |  |  |  |  |
| of growth, | **neg** | 582 | 31.6 | 4 | 11.8 | 122 | 32.2 | 3 | 16.7 |  |  |
| stemness | **pos** | 1259 | 68.4 | 30 | 88.2 | 257 | 67.8 | 15 | 83.3 |  |  |
| and EMT | **total** | 1841 |  |  |  |  |  |  |  | **0.011 (F)** | 0.202 |

## Discussion

In our previous report on PCa, BRCA1 gene losses were found to correlate with more advanced stage of the disease, dissemination of tumor cells and worse clinical outcome [1]. The present study was designed in order to validate these findings in a large cohort of patients. Furthermore, this study aimed to determine tumor molecular features associated with the aberrations of BRCA1 gene and potentially involved in PCa progression such as proliferation, epithelial-mesenchymal transition or stem-cell-like phenotype.

The distribution of BRCA1 gene aberrations was similar to the previously reported: BRCA1 gene losses appeared in 10% and gains in 7% of the analyzed patients [1]. The intratumoral heterogeneity was defined in almost one seventh of the tumors analyzed in two tumor samples, which confirms the molecular multiplicity of prostate cancer. Common molecular alterations of BRCA1 gene such as hypermethylation of the promoter and/or known germline base (ex)change mutations did not account for BRCA1 gene loss-of-function in tumors with BRCA1 gene gain. However, other chromosomal aberrations might be the causative that did not disrupt FISH probe binding but e.g. interactions with one of the multiple binding proteins of BRCA1 [20, 21]. Of note, 40% of the investigated tumors with BRCA1 gene gain were characterized by cytoplasmic BRCA1 protein expression or loss of nuclear BRCA1 expression. BRCA1 absence in nuclei and/or mislocalisation in cytoplasm might result in dysfunction of BRCA1 as tumor suppressor e.g. by affecting its role in DNA damage repair [21, 22]. Therefore, it might be assumed that at least a part of those tumors do not have fully functional BRCA1 protein, which merits further investigation.

The current study corroborated the associations of BRCA1 gene losses to more advanced tumor disease defined by higher T status and Gleason score in the large cohort of patients. BRCA1 gene losses indicated also shorter time to biochemical recurrence in the patients characterized by Gleason score >7. Of note, BRCA1 gene losses occurred to be an independent prognostic marker in this group of patients, which hypothetically might facilitate the treatment decisions in this high risk group of patients in the future. It is conceivable that such patients might profit from poly(ADP-ribose)polymerase (PARP) inhibitors already tested in some clinical trials or other sort of therapeutics expected to function in BRCA1-deficient tumors [22]. In addition, beside gene losses BRCA1 gene gains were detected, which also correlated to shorter time to metastasis.

Thirty-six percent of patients analyzed for matched tumor – LNM samples had BRCA1 gene losses in LNMs. Our previous results [1,17] and study on PCa metastases using next generation sequencing [4] suggest that not only germline mutations but also somatic BRCA1 gene losses might be associated with tumor dissemination to distant organs. In the current study, BRCA1 gene losses could have been tracked in both primary and secondary tumor site only in 2 patients. However, BRCA1 gene losses might be present only in one even smaller tumor focus [1,17] or in a small subpopulation of tumor cells as we have already shown by analysing whole sections of tumors [1]. It might be reasoned that patients with BRCA1 gene loss found exclusively in LNM might still carry BRCA1 gene losses in another focus of the primary tumor which had been missed during evaluation of small tumor fragments such as TMA tissue cores.

BRCA1 plays different roles including control of cell cycle [6] and speculated regulation of differentiation of breast (cancer) stem cells [23–25]. Herein, we cannot define a clear phenotype of PCa with BRCA1 gene losses or gains. Both tumors with BRCA1 gene losses and BRCA1 gene gains displayed more frequently ALDH1 known to characterize stem cells in different types of organs [26]. BRCA1 gene losses were also associated to the expression of EGFR, a regulator of plethora of different cellular processes including growth, stemness and EMT [27]. Based on these associations, it might be assumed that BRCA1 gene aberrations occur in the subpopulation of PCa stem cells or induce stem cell programme (dedifferentiation process) in PCa cells.

## Conclusions

In conclusion, in the current large study it was substantiated that BRCA1 gene losses present even in a small subpopulation of PCa might indicate a more aggressive stage of disease. Additionally, it shows that also BRCA1 gene gain marks more invasive tumors. Both types of BRCA1 gene aberrations are associated with ALDH1-expression, which would suggest a stem-cell like phenotype. In the future identification of BRCA1 gene aberrations might help to identify the patients with the highest risk of metastasis and even choose an appropriate therapy e.g. with PARP inhibitors.

### Ethics approval and consent to participate

This study complies with the Declaration of Helsinki. The samples and data collection was approved by local Ethics Committee (Ethik Kommission der Arzt kammer Westfalen-Lipp und der Medizinischen Fakultat der Wesfalischen Wilhelms-Universitat, nr 2007-467-f-S). The participants gave written informed consent for data collection and storage, as well as individual follow-up. All data were anonymized and treated with the highest caution.

## Availability of data and material

The datasets used and/or analysed during the current study are available from the corresponding author on reasonable request.

## Authors’ contributions

The authors’ responsibilities were as follows. NBK and BBr conceived and designed the research. AO, PN, AB, MD, BBi and NBK performed experiments and analysed the data. SH, AS, EE, BBr and KP provided the samples, original data and resources. PN, BBr, AS and KP advised on the study design, analysis and interpretation of the results. NBK was responsible for drafting the manuscript, whereas all of the authors read and approved the final manuscript. None of the authors had financial or non-financial interests relevant to the submitted manuscript.

## Supplementary Tables and Data

**Supplementary Table 1.** Separation conditions used for each exon of the BRCA1 gene.

| Exon | Mutation | dHPLC temp. (°C) | Gradient (%B) |
| --- | --- | --- | --- |
| 2 | 185delAG | 49 | 53.3 |
| 5 | Cys61Gly | 54 | 55.2 |
| 11 | 3819del5 | 53 | 55.0 |
| 11 | 4153delA | 56 | 57.7 |
| 20 | 5382insC | 58 | 55.1 |
| 20 | Arg1751Ter | 58 | 55.1 |

**Supplementary Table 2.** Comparison of BRCA1 gene dosage and protein expression. Comparison of BRCA1 gene dosage and protein expression in subset of PCa (n=203)

| BRCA1<br>gene<br>dosage | BRCA1 protein expression |  |  |  |  |  |  |  |
| --- | --- | --- | --- | --- | --- | --- | --- | --- |
|  | nuclear (+)<br>cytoplasmic (-) |  | nuclear (+)<br>cytoplasmic (+) |  | nuclear (-)<br>cytoplasmic (+) |  | nuclear (-)<br>cytoplasmic (-) |  |
|  | n | % | n | % | n | % | n | % |
| loss | 12 | 48.0 | 2 | 8.0 | 4 | 16.0 | 7 | 28.0 |
| normal | 79 | 34.5 | 21 | 12.5 | 10 | 6.0 | 58 | 47.0 |
| gain | 6 | 60.0 | 2 | 20.0 | 0 | 0 | 2 | 20.0 |

**Supplementary Table 3.** BRCA1 gene dosage in multifocal prostate cancer. Comparison of BRCA1 gene status in two different fragments of primary tumor (n=34).

| BRCA1 gene dosage in<br>tissue core nr 1 | BRCA1 gene dosage in<br>tissue core nr 2 | n | % |
| --- | --- | --- | --- |
| loss | normal | 3 | 8.8% |
| loss | gain | 0 | 0 |
| normal | loss | 9 | 26.5% |
| normal | gain | 9 | 26.5% |
| gain | loss | 1 | 2.9% |
| gain | normal | 12 | 35.3% |

**Supplementary Table 4.**
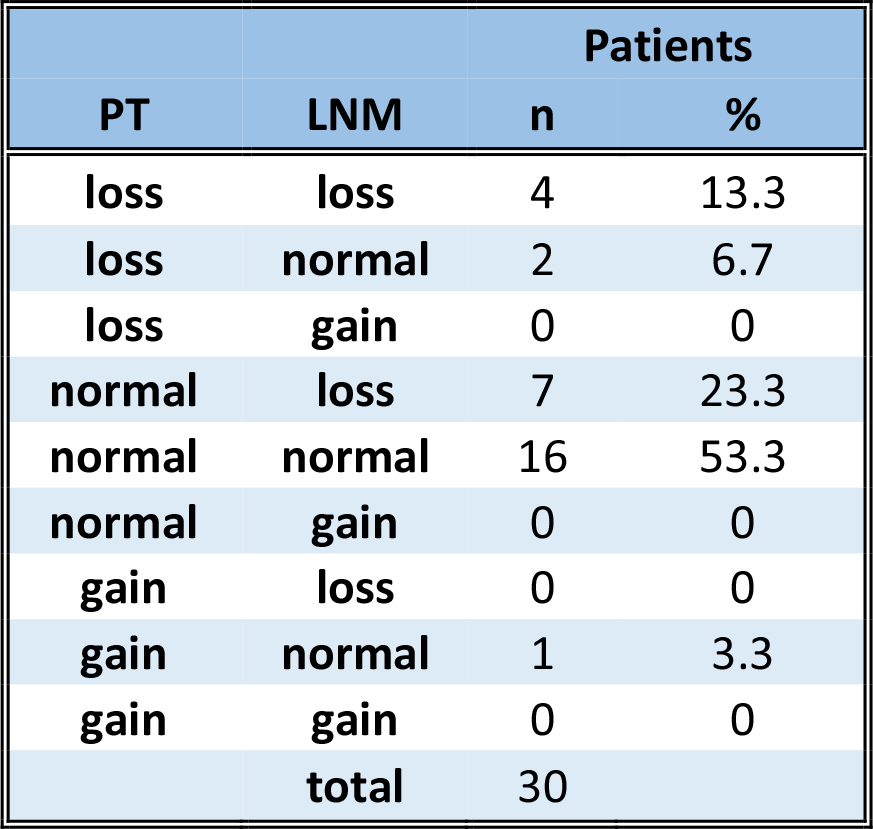
BRCA1 gene dosage in primary tumor– lymph node metastasis pairs. Comparison of BRCA1 gene status in matched pairs: primary tumor (PT) – lymph node metastasis (LNM).

## Supplementary Data

### Immunohistochemistry against cytokeratins

TMAs were preheated in 0.01M citrate buffer (pH 6.0, Dako, Denmark). IHC against cytokeratins (CKs) was performed with the usage of following monoclonal mouse antibodies (Ab): anti-CK5/6 Ab (clone D5/16B4, Dako, Denmark), anti-CK14 Ab (clone LL002, Dianova, Germany), anti-CK8/18 Ab (clone K8.8/DC10, Dianova, Germany) and anti-CK19 Ab (clone KS19.1, Quartett, Germany) diluted 1:40, 1:50, 1:40 and 1:500, respectively, incubated for 25 min. at room temperature. Stainings were envisioned with the usage of DAKO LSAB 2 System-AP (Dako, Cytomation, Denmark) and counterstained with haematoxylin.

### Immunohistochemistry against E-cadherin

Monoclonal mouse anti-E-cadherin Ab (clone NCH-38, Dako, Denmark) diluted 1:500 was incubated onto TMA section at room temperature after Tris-EDTA (pH 9.0)-based antigen retrieval in steamer. Staining was developed by LSAB/AP-Kit (Dako Cytomation, Denmark) in an automated system (DAKO Autostainer, Denmark) and counterstained with heamtoxylin.

### Immunohistochemistry against N-cadherin

IHC against N-cadherin was done after antigen retrieval performed in 0.01M citrate buffer (pH 6.0, Dako, Denmark) in steamer. Monoclonal mouse anti-N-cadherin Ab (clone 6G11, Dako, Denmark) was diluted 1:100 and incubated at room temperature. Staining was developed by LSAB/AP-Kit (Dako Cytomation, Denmark) in an automated system (DAKO Autostainer, Denmark) and counterstained with haematoxylin.

### Immunohistochemistry against vimentin

Monoclonal mouse anti-vimentin Ab (clone RV202, BD Pharmingen, USA) diluted 1:500 was incubated at 4°C for 16 hours on TMA section preheated in 0.01M citrate buffer (pH 6.0, Dako, Denmark) at 120°C for 5 min. Staining was envisioned by DAKO ChemMate Detection Kit Peroxidase/DAB, Rabbit/Mouse (Dako, Denmark) and counterstained with haematoxylin.

### Immunohistochemistry against Ki-67

Monoclonal mouse anti-Ki-67 Ab (clone MIB1, Dako Diagnostika, Germany) was applied in the dilution 1:1000 after antigen-retrieval in the pressure cooker at 121°C for 5 min. Staining was detected by LSAB/AP-Kit (Dako Cytomation, Denmark) in an automated system (DAKO Autostainer, Denmark) and counterstained with haematoxylin.

### Immunohistochemistry against apoptosis marker

Signatures of apoptosis were assessed by apoptosis detection kit ApopTags (Chemikon, Germany) according to the manufacturer’s protocol with minor modifications (pressure cooker at 37°C for 15 min.).

### Immunohistochemistry for ALDH1

Deparaffinised TMA sections were boiled for 5 min. in citrate buffer (pH 6.0, Biogenex, USA) at 120°C at steamer and incubated for 16 hours at 4°C with mouse monoclonal anti-ALDH1 antibody (44/ALDH1, BD Biosciences, US) diluted 1:500 in Dako REAL™ Antibody Diluent (Dako, Denmark). Staining was envisioned by DAKO ChemMate Detection Kit Peroxidase/DAB, Rabbit/Mouse (Dako, Denemark) and counterstained with hematoxylin (Merck, Germany).

### Immunohistochemistry for EGFR

Deparaffinized TMA sections were treated for 6 min. with Proteinase K Ready-to-Use (Dako) and for 5 min. with Perioxidase-Blocking Solution (Dako). TMAs were incubated overnight at 4°C with mouse monoclonal anti-EGFR antibody (E30, Dako) diluted 1:20 and envisioned by EnVision Kit, Rabbit/Mouse (Dako) and counterstained with hematoxylin (Merck, Germany).

### Evaluation of immunohistochemistry

Proteins’ expression was determined in 0 to 3 scale as follows: ‘0’ - no expression, ‘1’ – weak expression, ‘2’ – moderate expression and ‘3’ – strong expression. For further statistical analysis these values were analysed grouped into ‘0-1’ vs. ‘2-3’ classes as negative vs. positive expression, respectively. Additionally, for vimentin, Ki-67, apoptosis marker, ALDH1 and EGFR stainings percentage of marker-positive tumor cells was documented.

## Abbreviations

ALDH1: aldehyde dehydrogenase
CK: cytokeratin
CTCs: circulating tumor cells
DFS: disease-free survival
EGFR: epidermal growth factor receptor
EMT: epithelial-mesenchymal transition
FISH: fluorescent in situ hybridization
IHC: immunohistochemistry
LNM: lymph node metastasis
PARP: poly(ADP-ribose) polymerase
PCa: prostate cancer
PSA: prostate specific antigen
PT: primary tumor
REMARK: REporting recommendations for tumour MARKer prognostic studies
TMA: tissue microarray

